# Assisted gene flow yields *Acropora palmata* corals with robust physiological performance under warmer water temperatures in a land-based nursery

**DOI:** 10.1101/2025.03.03.641242

**Authors:** Erinn M. Muller, Chelsea Petrik, Trinity Conn, C. Cornelia Osborne, Marina Villoch, Abigail S. Clark, Hanna R. Koch, Keri O’Neil, Cody Engelsma, Iliana B. Baums

## Abstract

Assisted gene flow (AGF) is a conservation approach that facilitates the spread of alleles and may accelerate the recovery of genetically depauperate cohorts. The threatened Caribbean coral *Acropora palmata* is approaching regional extinction within the western Atlantic partly due to increasing water temperatures associated with global climate change. Previously, AGF was conducted by crossing gametes collected from three regions (Curaçao - CU, Florida - FL, and Puerto Rico - PR) characterized by contrasting temperature regimes and low gene flow between them. Here, we tested the thermal tolerance of these AGF cohorts in comparison to purebred Florida and Curaçao cohorts. Exposure to high temperatures resulted in few physiological changes, likely because the corals hosted the thermally tolerant algal symbiont, *Durusdinium trenchii*. However, the FL x FL cohort was the most sensitive to the high temperatures with a significant reduction in net photosynthesis and maximum electron transport rate under this treatment. Like the phenotypic responses, gene expression changes in response to heat stress were muted overall. Consequently, there was little power to detect correlations between genotype and phenotype. Relative to mid-parent values, CUxFL AGF cohorts showed 26 overexpressed and 48 underexpressed genes. Differentially expressed genes included known stress responders. Importantly, hybrid crosses harbored 879 private alleles that were previously not recovered in representative genets from Florida and thus carry important conservation value. These findings suggest that AGF corals not only carry novel alleles but also represent novel gene expression patterns.

## Introduction

Assisted gene flow (AGF) is the intentional translocation of individuals or gametes within a species’ range to transfer genetic diversity and facilitate adaptation to anticipated local conditions (Aitken and Whitlock 2013). With the rate at which anthropogenically induced climate change is occurring (IPCC 2018), bold interventions such as AGF may be needed to ensure the persistence of species that are unable to adapt to the rapid increases in global temperatures. AGF has already been employed for some forest trees (Morgenstern *et al*. 1990, Rehfeldt et al. 2001, Girardin *et al*. 2021) and had positive fitness outcomes across diverse animal taxa (Clarke et al. 2024). AGF is advantageous because the strategy can be implemented today rather than an approach such as gene editing, which is currently not possible for many traits and species on the brink of extinction. Indeed, AGF may be a critical tool for preserving species at risk of local extinction because of climate change, especially species that are at risk of inbreeding depression from recent large population declines. This is because AGF provides recipient populations with genetic diversity from elsewhere in the range and decreases the average kinship of offspring resulting from inter-population crosses. Increasing genetic diversity and decreasing average kinship are two critical population management goals that can be achieved via AGF.

Coral reefs are facing unparalleled levels of rapid ocean warming, which has led to an increase in the frequency and severity of coral bleaching events and the subsequent mortality of corals worldwide (Hughes *et al*. 2018a; Hughes *et al*. 2018b; Hughes *et al*. 2018c). The loss of corals has led to increased investments in coral reef conservation initiatives and the development of innovative restoration interventions for preserving species and reef function (van Oppen et al. 2015, 2017; NASEM 2018; Baums *et al*. 2019). In the eastern Atlantic and Caribbean region, the focus has been on the *Acropora palmata* and *A. cervicornis*, that have undergone extensive range-wide declines from disease outbreaks and bleaching events (Aronson and Precht 2001; Patterson *et al*. 2002; Williams *et al*. 2008, 2020). These species are listed as threatened under the US Endangered Species Act, critically endangered under the IUCN Red List, and *A. palmata* is currently facing regional extinction within Florida’s Coral Reef Tract (Hogarth 2006; Aronson et al 2008; Carpenter et al 2008, Williams et al. 2024).

*Acropora palmata* has been targeted in cryopreservation studies (Hagedorn *et al*. 2012, 2018a, 2021). Cryopreservation is an important conservation tool that has proven successful at preserving mammalian embryos (Fahy *et al*. 1984; Pollard and Leibo 1994; Kasai 1996), fish spermatogonial cells (Hagedorn *et al*. 2018b), and coral larvae (Daly *et al*. 2018), sperm (Hagedorn *et al*. 2006, 2012, 2017, 2018a, 2021), and microfragments (Powell-Palm et al. 2023). Since *A. palmata* gametes are only viable for a few hours post-release, cryopreservation of the sperm has tremendous implications for the propagation and restoration of *A. palmata*. To determine if cryopreservation could be used to facilitate AGF among geographically separated and genetically isolated populations of *A. palmata*, Hagedorn et al. (2018) collected and cryopreserved sperm from *A. palmata* colonies located in the western Atlantic (Florida, USA), central Caribbean (Puerto Rico), and southern Caribbean (Curaçao). The cryopreserved sperm was then crossed with freshly collected eggs from Curaçao to produce viable offspring, demonstrating that cryopreservation can facilitate AGF among *A. palmata* populations and potentially play an important role in the conservation of population-level genetic diversity as well as improve population resilience.

Despite the intended benefits of AGF, unintended negative outcomes are possible including outbreeding depression which is the reduction in fitness resulting from mating genetically distinct populations (Frankham *et al*. 2011). Potential mechanisms include the loss of locally adapted alleles (Templeton *et al*. 1986), or disruption of co-adapted genes at different loci (epistasis) (Edmands 2007). Outbreeding depression is possible between Florida and Curaçao populations insofar as these locations may not be regularly exchanging genetic diversity (Baums *et al*. 2005; Baums *et al*. 2006a, 2006b; Vollmer and Palumbi 2007). Puerto Rico, however, has been identified as an area of mixing between the western and eastern Caribbean regions (Baums *et al*. 2005; Baums *et al*. 2006a). Even though outbreeding depression has never been directly tested in stony corals, associated risks are predicted to be low (Baums *et al*. 2019), compared to its potential benefits in rapidly declining local populations that are failing to reproduce sexually (Ralls *et al*. 2018), such as *A. palmata* in the Florida Keys.

Another consideration of performing AGF with stony corals is that these species possess an obligate, mutually beneficial symbiosis with unicellular algae (Family: Symbiodiniaceae) that can influence host physiology and stress responses (Mieog *et al*. 2009; Howells *et al*. 2012). Therefore, the phenotype of an adult coral is the result of the algal endosymbiont and host together (Parkinson and Baums 2014) (Parkinson and Baums 2014). This coral-algal symbiosis is especially important when it comes to thermal tolerance and coral bleaching. Under prolonged temperature extremes, the symbiosis breaks down resulting in endosymbiont expulsion, coral ‘bleaching’, starvation and eventually death if the stress does not subside and symbiosis is re-established. However, symbiont species differ in their physiological optima and influence the stress phenotype of their host under changing environmental conditions (Little *et al*. 2004; Stat and Gates 2011; Cunning *et al*. 2015). *A. palmata* associates with *Symbiodinium ‘fitti’* (nominem nudum) and adults only rarely host other symbionts such as *Durisdinium trenchii* or *Cladocopium* sp. (Baums et al. 2014). Juvenile *A. palmata* are more promiscuous and sometimes host *D. trenchii*, which could be retained for years in adults (Elder et al. 2023). Because *D. trenchii* imparts a measure of thermal tolerance to its hosts, we established the symbiont species infecting AGF corals and investigated host and symbiont traits.

Within the present study we assessed the physiological responses of the coral symbiont, host gene expression, and host population genetic structure of F1 offspring characterized as assisted gene flow (Curacao x Puerto Rico and Curacao x Florida) or pure-bred (Florida x Florida and Curacao x Curacao) cohorts. This study aimed to compare the following responses under non-stressful ambient water temperatures and prolonged heat stress conditions: coral metabolism (photosynthesis (P) and respiration (R) rates, daily P:R ratios), growth, a suite of algal symbiont photophysiology traits, gene expression profiles using RNASeq, and population genetic structure using single nucleotide polymorphism (SNP) loci.

## Materials and Methods

### Study site, species, and cohorts

The present study was conducted at Mote Marine Laboratory’s Elizabeth Moore International Center for Coral Reef Research & Restoration facility (24°39’41”N, 81° 27’ 16” W, Summerland Key, Florida, USA). The *Acropora palmata* corals used in this study originated from Hagedorn *et al*. (2021), which were subsequently maintained in Mote’s land-based nursery and The Florida Aquarium. The corals within the present study represent four different *A. palmata* cohorts (crosses): i) a batch cross of fresh eggs and sperm from the Upper Florida Keys (FL x FL); ii) a batch cross of fresh eggs and sperm from Curaçao (CU × CU); iii) a cross between fresh eggs from Curaçao and cryopreserved sperm from Florida (CU × FL); and iv) a cross between fresh eggs from Curaçao and cryopreserved sperm from Puerto Rico (CU × PR). The CU x FL and CU x PR crosses represent the ‘AGF’ hybrid cohorts, while the other two represent ‘pure-bred’ cohorts. For population-level analyses, we further included existing genotyping data from randomly selected wild Florida Keys (n=30), Curaçao (n=30) and western Puerto Rico (n=27) genets. These genets were genotyped using the same methods as the experimental crosses and are publicly available in the STAG database (coralsnp.science.psu.edu/galaxy). Genotyping methods are described below.

### Experimental set-up

The thermal stress experiment was carried out within Mote’s Climate and Acidification Ocean Simulator (CAOS) system from 15 June to 2 September 2020. A total of 50 putatively different genotypes from each of the four cohorts reared within Mote’s land-based coral nursery were initially fragmented in half. Colonies were fragmented to 1-2 cm^2^ in size using a band saw (C-40 diamond Band saw, Gryphon) and mounted onto flat ceramic plugs (1 ¼” diameter, Boston Aqua Farms) with cyanoacrylate gel. The fragmented corals were given two weeks to recover and acclimate within the CAOS system before the onset of the experiment. One of each replicate per cohort (n=50) was placed within the ambient temperature treatment (control) and the other replicate was placed within the high temperature treatment (experimental). This created a total of 200 fragments per treatment (50 replicates per cohort, four cohorts per treatment).

To assess cohort-level responses to heat stress, coral fragments were subjected to one of two temperature treatments, ambient (28°C) or elevated (31°C), for two months. Ambient temperature was modeled after the mean temperature for June curated from temperature loggers at Looe Key Reef, which has recorded in situ temperature at 6 m for the past 15 years. The temperature of 31°C was selected for the elevated temperature treatment as 30.5 °C is estimated to be the bleaching threshold reported for Acroporid corals in the Florida Keys (Manzello *et al*. 2007, Williams et al. 2017). On 8 June 2020, mounted corals were placed within their respective 20 L flow-through glass tanks (16.25” x 8.375” x 10.5”) fitted with a transparent Plexiglas lid and submersible pumps (120 GPH; Dwyer). Forty tanks were held within four shallow fiberglass flow-through mesocosms (‘raceways’)(100” x 40” x 12”) under an overhead shaded structure (∼75% light reduction) outside. Each tank held 9-10 replicates from a single cohort and each cohort consisted of 5 separate tanks per treatment. Temperature was regulated in the raceway and maintained at either ambient (28°C) or high (31°C) conditions. On 25 June 2020 (Figure 1), after preliminary physiological measurements were performed, temperatures were manually raised in the water baths containing respective high temperature treated experimental tanks at a rate of +0.5 °C per day until the target temperature of 31 °C was met. Temperatures were controlled by a digital temperature controller (TSW-250, Dwyer) with inline sensors and heat exchangers via a central heater (Lochinvar-Knight, 286 MBH Hi-Efficiency Boiler) or chiller (15 Ton Air Chilled Cooler). Once at target temperature conditions, the temperatures were held constant (deadband ± 0.2 °C) for 8 weeks.

**Figure 1.**
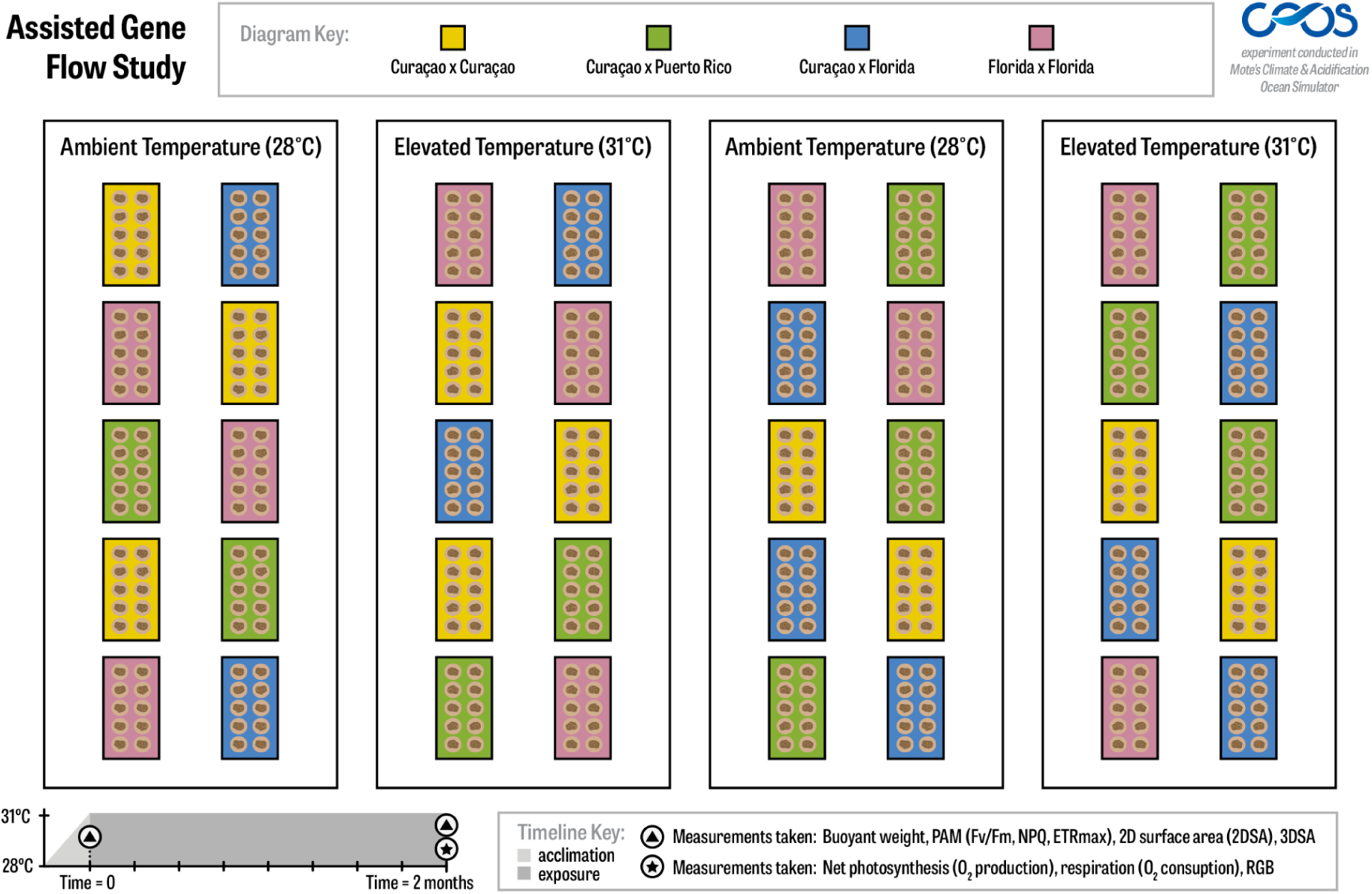
**Schematic representation of the experimental design used within the present study.**

Water quality parameters including temperature (°C), pH_NBS_ (NBS: National Bureau of Standards), dissolved oxygen (DO; mg/L), and salinity (ppt) were measured daily for each experimental tank using a multi-parameter handheld containing a conductivity/temperature sensor, galvanic DO sensor, and a pH sensor (YSI professional series sensors). The pH sensor was calibrated daily using NBS scale standard buffers (4, 7, and 10). Corals were fed once a week near dusk (100-200 microns, Golden Pearls, Aquatic Foods Inc.). Light levels (photosynthetically active radiation) were recorded with a planar underwater quantum sensor (LI-1500 and LI-192, Li-Cor) near maximum irradiance (approx. 13:00 GMT -5) every three days.

### Growth rates

Coral fragment calcification rates were measured using the buoyant weight method (adapted from Davies 1989). Corals were weighed prior to exposure and again after eight weeks of exposure by placing the coral on a suspended egg-crate (diffuse styrene lighting panel) platform connected to a monofilament line below a precision balance where the stabilized weight was recorded. One coral per tank was selected haphazardly to be weighed in duplicate to ensure accuracy of the scales; the difference between the duplicate weights was low (<0.008 g) suggesting dependable measurements. Dry weight was calculated from methods described by Jokiel et al. (1978) and the change of weight was characterized, as a relative unit, by subtracting the initial dry weight from the final dry weight and adjusted per initial two-dimensional (2D) surface area per day (cm^2^ day^-1^). During weighing periods, overhead photographs were taken of each coral replicate to derive 2D surface area (cm^2^) where image analysis was performed in ImageJ (National Institutes of Health). Change of 2D surface area was calculated from the difference between the final size and the initial size and adjusted per day (cm^2^ day^-1^).

The three-dimensional (3D) surface area for twenty replicates per cohort per temperature treatment was measured using a structured-light 3D scanner. Coral images were taken using a HDI Compact scanner (C210 Polyga, Burnaby, BC Canada) on a rotating plate following the methods of Koch et al. (2021). Images were captured every 30° in one rotation, from one angle, to produce 12 meshes. The High Dynamic range (HDR) setting was used as dual exposure was necessary to appropriately capture and differentiate between surfaces of the ceramic plug (coral mount) and actual coral tissue. The 3D scanner was calibrated prior to measurements using a 5 mm calibration board. For the analyses, meshes were uploaded to the Flexscan3D software to be reconstructed manually, and finalization of the model was performed. Mesh editing (i.e., hole filling and smooth merge functions) was applied where necessary. Once the 3D model was finalized, the model was trimmed to exclude the plug. Once editing and trimming were completed, the model was measured for remaining surface area (mm^2^) as described by Koch et al. (2021). Change in 3D surface area was calculated from the difference between the final size and the initial size (mm^2^).

### Respirometry

Twelve replicate corals per cohort per temperature treatment were measured for oxygen production and oxygen consumption at the end of the 8-week experiment. Net photosynthesis (P_net_) and respiration (R) rates were measured with a fiber-optic oxygen meter (Firesting O2, Pyro Science GmbH, Aachen, Germany) connected to a computer with Pyro Oxygen Logger software. An oxygen sensor was placed in each incubation chamber (300 mL) coupled with magnetic stir bars for adequate mixing. The oxygen sensors were calibrated prior to measurements in 100% air saturated water. Chambers were surrounded by a recirculating jacket at the appropriate treatment temperature (28 °C or 31 °C). Chambers were filled with seawater from the corals’ respective tank for every measurement. Each chamber contained one replicate coral. One chamber per each incubation period contained seawater only as an empty control, or ‘blank’, to correct rates for background microbial respiration. A standard fluorescent light strip was used as a lighting source and placed overhead at a fixed distance from the chambers. Photosynthesis rates (oxygen production) were measured at approximately 52 ± 11 µmol photons m^-2^ s^-1^ for 60 min. Respiration (oxygen consumption) rates were measured in complete darkness for 60 min. Rates were normalized per two-dimensional surface area (cm^2^) of the coral fragments taken at the final time point. Daily P:R was calculated following the equation in Krueger (2019), where daily P:R was the P_gross_ multiplied with the number of daylight hours (12 h) and then divided by the product of 24 h times the hourly respiration value.

### Algal symbiont traits

An Imaging pulse amplitude modulated (I-PAM) fluorometer (Walz) was used to measure photosynthetic efficiency of the algal endosymbiont, the maximum potential quantum efficiency of photosystem II (F_v_/F_m_). Instrument settings were as follows: light intensity=6, saturation pulse intensity=8, gain=1-3. Measurements were taken at a fixed distance from the coral tissue in the center of the fragment. Corals were placed in complete darkness for 30 min prior to each measurement. A weak measuring light (<1 µmol quanta m^-2^ s^-1^) was used to obtain the minimum fluorescence (F_o_) followed by a saturating pulse (3000 µmol quanta m^-2^ s^-1^) to measure the maximum fluorescence (F_m_), which were used to calculate the ratio of variable fluorescence to maximal fluorescence (F_v_/F_m_). F_v_/F_m_ was measured for all fragments prior to administration of experimental treatments and after 8 weeks. After F_v_/F_m_ measurement, light intensities ranging from 0-1280 µmol quanta m-2 s-1 on intervals of 20 sec were emitted on the sample to produce rapid light curves (RLCs). ETR*_max_* was determined by multiplying the effective quantum yield (measured during RLCs) by photo flux density (400-700 nm) (Platt et al. 1980). The maximum level of non-photochemical quenching within the light curve (NPQ*_max_*) was calculated as (Fm-Fm’)/Fm’; where quenched Fm is called Fm’ (Krause and Weis, 1991; Ruban 2016).

Overhead two-dimensional photographs were taken of each fragment at three time points (pre-exposure, one month post exposure, two months post exposure) to measure change in tissue surface area, as previously described, and change in pigmentation. Photographs were taken with a digital camera (Olympus TG-6) held at a fixed distance above the corals where a metric ruler and white/black color calibration card were placed within frame on the same plane as the coral fragment. Camera settings for all photographs were pre-fixed and included an aperture speed of 1/60s, ISO=120-160, F= 2.8, with the flash off. Color analysis was completed at the final time point based on the red, green, and blue (RGB) color model as described by Winters et al. (2009). RGB channel color intensity was based on the 0-255 scale. Normalization of image color was performed prior to a histogram analysis for each contributing color in GIMP software (version 2.10).

### Statistical analysis of phenotype data

Permutational multivariate analysis of variance (PERMANOVA, adonis function in the R package ‘vegan’ [Version 2.5-7, Oksanen *et al*. 2020]) were conducted to determine whether there were significant differences in the phenotypes among the four cohorts within the two temperature treatments (i.e., ambient and high). To visualize the data, principal components analyses (PCA; function prcomp in R base program) were applied to characterize the physiological response parameters among the four cohorts within each of the two temperature treatments. Traits that were significantly correlated were removed from the PCA. P:R was removed as it was correlated with P_net_ and R. Additionally, ETR*_max_*, F_v_/F_m_, and NPQ were all correlated so only F_v_/F_m_ was retained in the multivariate analysis. The contribution of each phenotype to the first two PCA axes were determined using the ‘get_eigenvalue’ and ‘get_pca_var’ functions within the package ‘factoextra’ (Kassambara and Mundt 2020). Data was normalized prior to the PERMANOVA and PCA applications to create comparable units.

To determine the effect of temperature treatment for each phenotype metric, individual Welch Two Sample t-tests were conducted with temperature treatment as the fixed factor on each physiological parameter. Normality was evaluated using a Shapiro-Wilk test and homoscedasticity of variances was assessed using Levene’s test (LeveneTest function, *car* package, Fox and Weisberg 2019). Where log transformations failed to meet parametric assumptions, Wilcoxon rank sum tests were used. Cohorts were analyzed separately to better assess the effects of temperature treatments on each cohort independently.

### Tissue sampling for gene expression and cohort genomic analyses

From 11-13 June 2020, all corals (n=500) were sampled to determine baseline gene expression levels before heat stress exposure. Using a sterile precision knife, two to five polyps were carefully removed from each coral and immediately placed in a 2 mL cryogenic tube containing 1.5 mL RNA*later* (Invitrogen), followed by storage in a 4 °C refrigerator for 24 hrs. After 24 hrs, the samples were transferred to a -80 °C freezer and stored until further analysis. Once the physiological measurements were completed (at the 8-week time point), all remaining corals were resampled for gene expression analysis (n=420) and for population genomic analysis (n=426) post-heat stress. Thirty-six corals had significant tissue loss prior to the end of the experiment (5 September 2020) and were therefore sampled before the conclusion of the study. An additional 36 corals suffered complete mortality during the experiment and were consequently not sampled. In total, 456 corals were sampled for gene expression and population genomic analyses. After samples were collected for gene expression analysis, all remaining tissue was flash-frozen and stored at - 80 °C. Samples were transported to Pennsylvania State University and stored at -80 °C until further analysis.

### DNA extraction and genotyping

Samples were collected using a razor blade from all 250 genets included in the experiment. Samples were extracted using the QIAGEN DNeasy Blood and Tissue kit following published protocols (Kitchen et al., 2020). DNA was quantified at ThermoFisher via picogreen, and DNA quantities ranged from 1.71 to 79.38 ng µl^-1^. DNA was sent to ThermoFisher for genotyping using the Applied Biosystems Axiom Coral Genotyping Array – 550962 following Kitchen et al. 2020. Quality control analyses were performed by Thermofisher. 19,969 probes on the array were designed to resolve the population genomic structure of *A.palmata* and *A.cervicornis.* Subsequent analyses were performed using this set. After removing all samples that failed or were not included in the physiological measurements, 191 samples remained for subsequent analyses.

### Symbiont Typing

Of the 4,021 probes designed for the Symbiodineaceae hosted by coral, 54 probes were designed specifically to resolve genera (Kitchen et al., 2020). The raw signal intensities of these probes were exported using the Axiom Analysis Suite and using methods described in Kitchen et al., 2020, a Linear Discriminant Analysis (LDA) was used to determine the genera of Symbiodiniaceae was hosted by each sample of *Acropora palmata*.

### Genetic Diversity Analyses

To quantify genetic diversity across all four AGF cohorts and three wild populations, a Principal Component Analysis (PCA) was performed on genotyped samples using all 19,965 SNPs (single nucleotide polymorphisms). Next, data from wild genets from the Florida Keys (n=21), Curaçao (n=27), and Western Puerto Rico (n=35) was randomly selected from the Acropora StagDB (www.coral.snp.psu.edu) and added to the AGF cohorts to quantify how genetically similar the AGF cohorts were to their source populations (Figure 7b.)

Pairwise kinship coefficients were calculated for all corals using the KING robust method (Manichaikul et al. 2010). The experimental corals were bred from a small pool of parents. Therefore, we expected that experimental corals would be more closely related to each other than a comparable number of wild colonies from any particular reef. This is of interest because closely related genets are more likely to respond to stressors similarly than unrelated genets (Dixon et al. 2015, Howells et al. 2021). An ANOVA was performed to calculate the differences in genetic similarity within and between cohorts. To assess whether these corals were representative of their wild source populations, allelic diversity was quantified using the package PopGenReport (Adamack & Gruber, 2014) in R (R Core Team). Allelic richness was quantified across all markers and an ANOVA was performed to compare allelic richness across the genome among all cohorts. Private alleles had a frequency of 0.01 to 1.0 in the representative wild genets. We then determined the number of alleles carried by the AGF hybrids not found in the respective natal populations using a custom R script. Observed heterozygosity, expected heterozygosity, and FIS (inbreeding coefficient) were calculated for each cohort using the R package dartR (Gruber et al., 2018). All statistical analyses were performed using R v.4.0.2.

### Genotype-Phenotype Association

To test if there were any genotyped SNPs associated with the observed differences across cohorts, a genome-wide association study (GWAS) was performed using plink2 (Chang et al., 2015). The association test was performed on 5 traits with large enough sample sizes: 2D Surface Area, Buoyant Weight, F_v_/F_m_, NPQ_max_, and ETR_max_. Of 19,965 SNPs, 6445 polymorphic SNPs were retained for analysis. 377 of those SNPs were removed due to a minor allele frequency below 0.01. A KING kinship cutoff of 0.2 was established to prune any closely related recruits. To account for population structure, the top 10 principal components of the population structure were extracted using plink2 and input into the linear model as a covariate. Traits were analyzed as the change in each trait between the first sampling time-point and the final sampling time-point in all elevated-temperature samples. A Bonferroni correction was applied to determine alleles that showed a significant association with the tested phenotypes.

### RNA extraction and differential gene expression analysis

Of the 956 samples collected for gene expression, total RNA was extracted from samples that had both pre- and post-sample time points (*n=*810). For each sample, 2-3 polyps were homogenized in 1 ml TRIzol reagent (Ambion, Life Technology, USA) before centrifugation with 200ul chloroform for 15 minutes at 12,000 x g, 4 °C. The aqueous phase was removed and cleaned using Qiagen RNeasy Mini Kit, as per manufacturer’s protocol with an additional on-column DNase treatment (RNase-Free DNase Set, Qiagen). The eluted RNA (in RNAse-free water) was twice passed through the spin column to maximize final concentration. Concentration and purity (A260/280, 260/230) were analyzed via NanoDrop ND-1000 spectrometer and quality assessed (RIN > 7) with Agilent 2100 Bioanalyzer (Agilent Technologies). 498 high quality samples were submitted to Admera Health (New Jersey, USA) for removal of salt and 2×150bp paired-end sequencing using the NEBNext Ultra II Nondirectional Library Prep Kit with polyA selection (New England Biolabs, Inc.) Illumina universal adapters and reads below PHRED of 23 were trimmed using Cutadapt (v.3.5, Martin 2011). Filtered reads were mapped to the *Acropora palmata* genome (v3.1, Kitchen 2020, unpublished) using *STAR* (v2.7.0a) with read count data generated by the -quantMode GeneCount parameter. Samples with alignment below 45% were excluded from further analyses, leaving 494 samples for DESEQ2 input.

Differentially expressed genes (DEGs) were identified with the R package *DESeq2* (Love et al., 2014). Genes that had fewer than 10 counts across any gene were discarded, leaving a total of 17983 genes in the dataset. We first fit the model Expression ∼ Cohort on the baseline expression data to identify any cohort-specific transcriptomic differences that may differentiate the populations under ambient conditions. To then assess the transcriptomic response to elevated temperatures, the model Expression ∼ Cohort * Treatment was used and pairwise contrasts were extracted to analyze (1) the cohort-specific gene expression under heat stress and (2) the differential response between cohorts to heat stress. Genes identified as differentially expressed were corrected for multiple testing using Benjamini and Hochberg false discovery rate (FDR) (Benjamini & Hochberg, 1995). Over-representation tests of gene ontology (GO) terms were performed using the package *clusterProfiler* (Yu et al., 2012) using a custom GO database for the *Acropora palmata* genome created with the *AnnotationForge* package (Carlson & Pagès, 2024) courtesy of Kitchen et al., in prep. Using the variance stable transformation (vst) counts as input, the ‘plotPCA’ function from *DESeq2* was used for PCA analyses to explore cohort, genotype, and treatment effects as well as identify other potential sources of variation such as tank effects and differentiating physiological measurements identified in this study. Tank, runway, and PAR were all included in initial models but were not found to be significant factors explaining clustering of the cohorts and were thus excluded from the final model.

In addition, to investigate baseline expression deviation of the hybrid crosses from pure-bred crosses, the hybrid-cross samples of CUxFL were compared against the parental samples of FLxFL and CUxCU, fitting the model ‘expression ∼group’. For this analysis, we used median expression values from the two replicate fragments of each genet in the pre-exposure ambient and pre-exposure heat-stress treatments. Expression in the hybrid crosses was then categorized as either greater than both parent crosses or below both parent crosses. Using the command rlog() in DESeq2, gene counts were normalized and log-transformed before use in PCA and heatmaps.

## Results

### Exposure conditions

The water quality conditions within the experimental tanks remained consistent for the 8 weeks the temperature exposure experiment was conducted (Table 1). Salinity was maintained within a range of 37.4 to 37.5 ppt, dissolved oxygen from 5.81 to 6.00 mg/L, pH_NBS_ from 8.00 to 8.02, and light levels from 410 to 426 umol photons m^2^ s^-1^. The temperature target for the control conditions was 28 °C and the high temperature treatment target of 31 °C were met, the experiment tanks were held at 28.0 ± 0.01 °C (mean±SE) in the control temperature treatment tanks and held at 31.0 ± 0.01 °C in the high temperature treatment tanks across the 8-week exposure period.

**Table 1.** Average (±SE) water quality conditions (temperature, salinity, pH_NBS_, dissolved oxygen, and light) measured in experimental tanks containing the four cohorts held at ambient and high temperature treatments across the exposure period from 1 July 2020 to 4 September 2020.

| Temperature Treatment | Temperature (°C) | Salinity (ppt) | pH (NBS) | Dissolved Oxygen (mg/L) | Light (PAR) |
| --- | --- | --- | --- | --- | --- |
| Ambient | 28.0±0.01 | 37.4±0.04 | 8.02±0.01 | 6.0±0.0 | 410±10 |
| High | 31.0±0.01 | 37.5±0.04 | 8.0±0.002 | 5.8±0.02 | 426±9 |

### Physiological response

There were no significant differences detected among cohorts when the physiological data were analyzed using a multivariate approach. This held true under both the ambient and the high temperature conditions (PERMANOVA ambient: F=1.31, df=3,35, R^2^=0.100, p=0.28; Figure 2a; PERMANOVA high: F=0.929, df=3,32, R^2^=0.080, p=0.435, Figure 2b). Under ambient conditions dimension 1 of the PCA accounted for 32.7% of the variance in the ordination plot, with 3D surface area (3DSA) driving that distribution along the axis (33.03%). Similarly, the second dimension of the ordination accounted for 21.1% in the variation among the data and was primarily driven by Respiration (32.79%). Under high temperature conditions dimension 1 of the PCA accounted for 27.7% of the variance with F_v_/F_m_ accounting for much of that distribution (37.35%). Dimension 2 of the PCA under high temperature conditions accounted for 26.6% of the variation and was primarily driven by calcification rates (BW, 41.08%).

**Figure 2.**
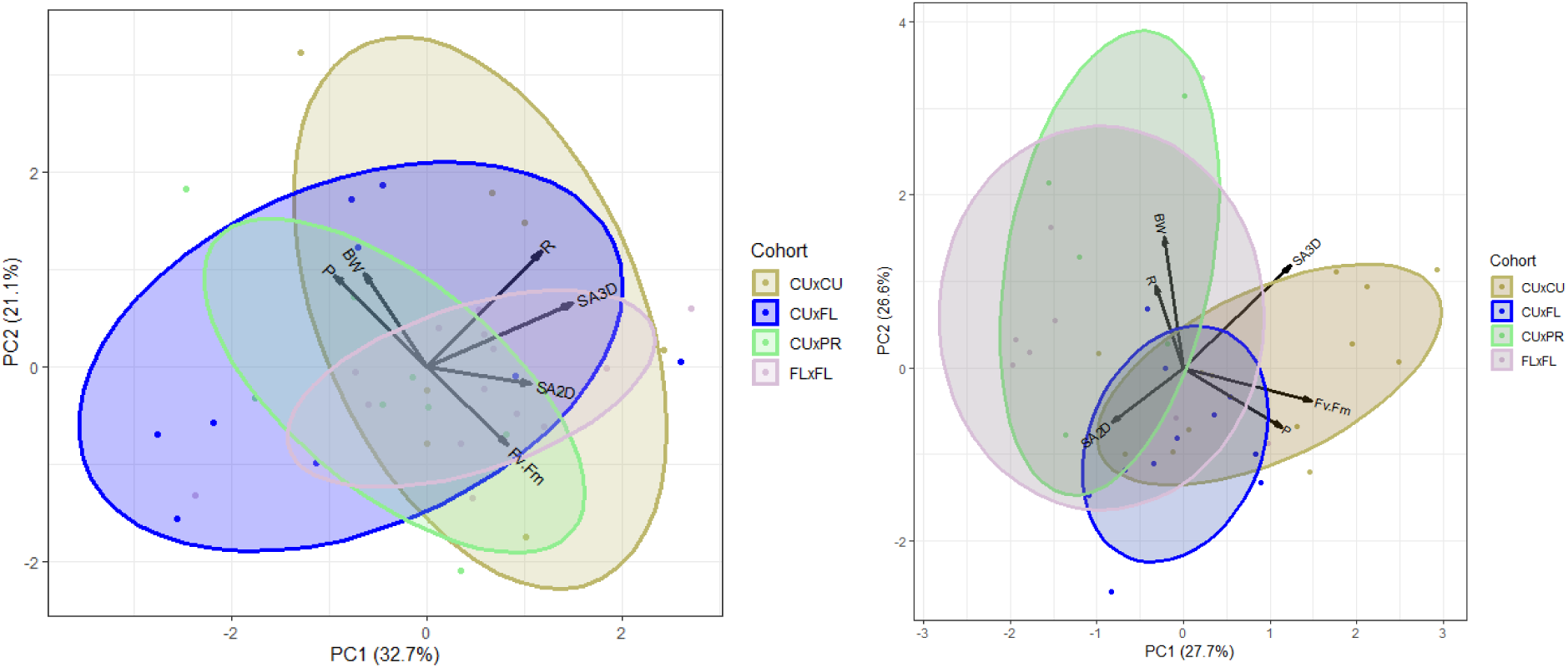
Distance biplots resulting from the principal component analyses (PCA). a) The response of four cohorts of *Acropora palmata* under ambient conditions. Coloration of the ellipses and points represents the four cohorts: yellow represents the Curaçao x Curaçao (CU x CU) cross, blue represents the Curaçao x Florida (CU x FL) cross, green represents the Curaçao x Puerto Rico (CU x PR) cross, and pink represents the Florida x Florida (FL x FL) cross. b) The overall response of four *Acropora palmata* cohorts under high temperature conditions. Physiological traits (black vectors) used to describe 2D surface area growth (SA2D), 3D surface area growth (SA3D), Calcification rates (BW), Respiration rates (R), Net Photosynthesis rates (P_net_), Photochemical efficiency metrics (F_v_/F_m_).

### Growth Metrics (2DSA, 3DSA, BW)

When the treatment temperature conditions were analyzed separately by cohort, there were no significant differences detected in 2D surface area (2DSA) between the control and high temperature treatments for three out of the four cohorts although all showed the trend of higher 2DSA growth under the high temperature treatment. One cohort, CU x PR, showed significantly higher 2DSA growth under the high temperature conditions compared with the control temperatures (t-test, t=-2.45, p-value = 0.02; Figure 3a). The 3D surface area (3DSA) growth rates of the corals from each cohort showed no significant differences between the two temperature treatments although all cohorts, again, showed a trend toward higher growth rates under the high temperature treatments (Figure 3b). There were no significant differences in calcification rate (BW) detected between the high temperature and control treatment within any cohort tested indicating they all accreted a similar amount of skeleton regardless of temperature conditions (Figure 3c).

**Figure 3.**
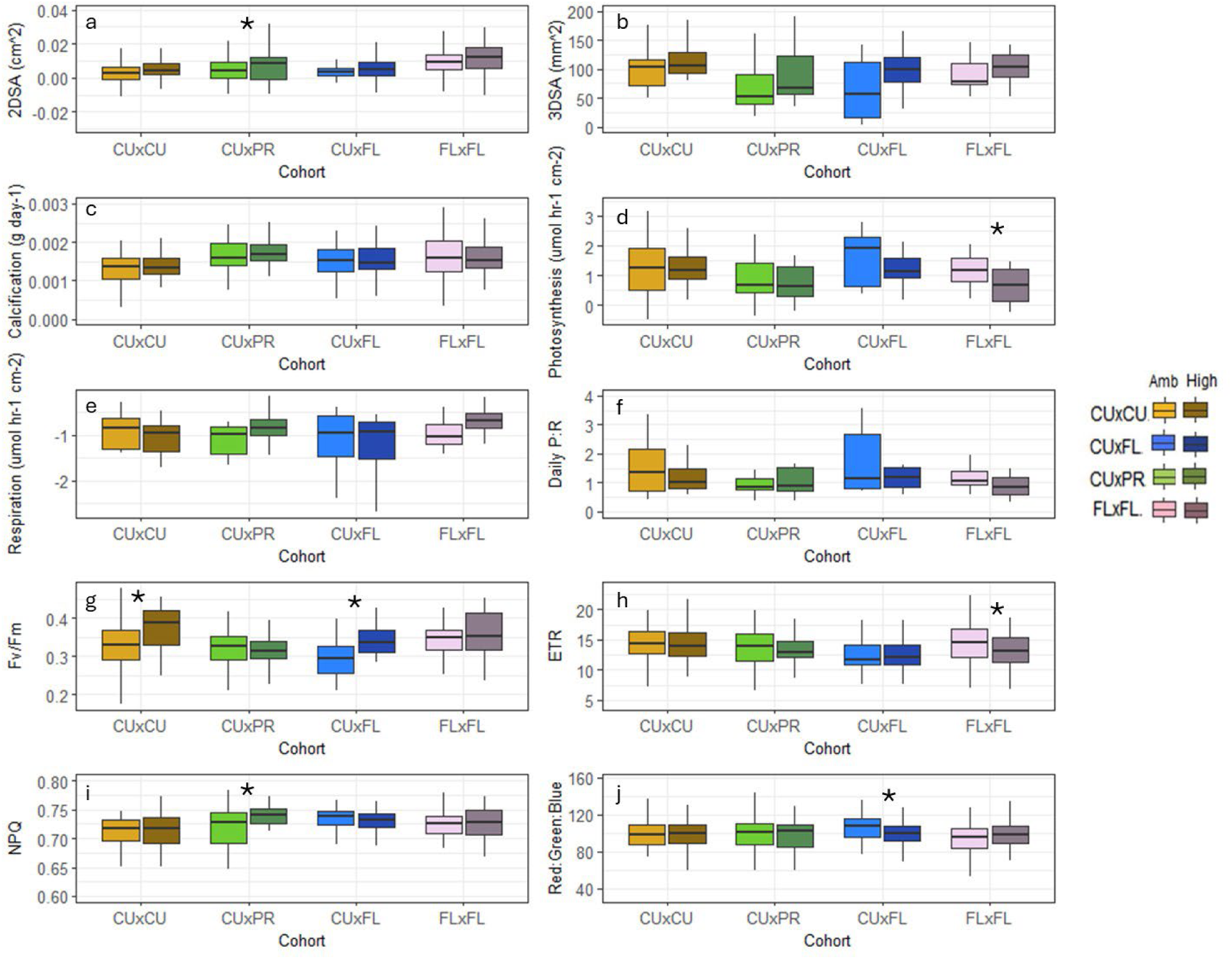
Effects of ambient (28°C) or high (31°C) temperature on four cohorts of *Acropora palmata* (Yellow: CU x CU, Green: CU x PR, Blue: CU x FL, Pink: FL x FL) on Change in 2D Surface Area (a), Change in 3D Surface Area (b), and Change in Buoyant Weight (c), Respiration rates (d), Photosynthesis rates I, P:R (f), F_v_/F_m_ (g), ETR_max_ (h), NPQ_max_(i), and Red:Green:Blue ratio (j). Boxplot whiskers show minima and maxima; centers indicate medians; box boundaries indicate the 25^th^ and 75^th^ percentiles. Asterisks denote significant differences between temperature treatments within a cohort.

### Respirometry (P_net_, R, daily P:R)

When each cohort was analyzed separately, three of the four cohorts showed no significant effect of temperature on net photosynthetic rates (P_net_) rates (Figure 3d). However, the FL x FL cohort exhibited significant decreases in their net photosynthetic rates under the high temperature treatment in comparison to the control temperature treatment (t-test, t=2.64, *p*=0.014; Figure 3d). FL x FL, CU x FL, CU x CU, and CU x PR had similar respiration rates regardless of temperature treatment (Figure 3e). When assessing the daily P:R ratio, there were no significant differences between the ambient and the high temperature treatments for any cohort (Figure 3f).

### Algal symbiont traits (F_v_/F_m_, ETR*_max_*, NPQ*max*, RGB)

The CU x PR and FL x FL cohorts showed no significant difference in F_v_/F_m_ values in the high temperature treatment compared with the control temperature treatments. However, the CU x CU cohort and the Cu x FL cohort had significantly higher F_v_/F_m_ values when held within the high temperature treatment compared with the ambient temperature (CU x CU: t-test,t=-2.72,*p*=0.008; CU x FL: t-test, t=-5.84 *p*< 0.001; Figure 3g). There was also a significant reduction in ETR*_max_* within the FL x FL cohort under high temperature conditions compared with ambient (t-test, t=2.26, p=0.026; Figure 3h). All other cohorts showed no differences in ETR*_max_* due to treatment conditions. Interestingly, the CU x PR cohort had a significant increase in NPQ*_max_* under the high temperature treatment (Wilcoxon test, W=632, *p*=0.001; Figure 3i). Finally, there was a significant decrease in the RGB values (i.e., higher algal symbiont concentrations) for the CU x FL cohort under high temperature conditions compared with ambient conditions (t-test, t=2.34, p=0.021; Figure 3j).

## Gene Expression Results

### Baseline Transcriptomic Comparisons

Gene expression analyses at baseline conditions revealed some variation among the coral cohorts, as visualized by principal component analysis (PCA, Figure 4*a*) and quantified through differential gene expression. The hybrid Curaçao x Puerto Rico (CU x PR) crosses exhibited the most differentiated transcriptomic profile, clustering distinctly from other cohorts along the first principal component (PC1), which explained 25% of the variance between cohorts. Comparisons between CU x PR and cohorts with a Floridian genetic background yielded the highest number of differentially expressed genes (DEGs) with 3,928 and 4,104 DEGs found with FL x FL crosses and CU x FL crosses, respectively. Gene Ontological (GO) Enrichment analyses showed pathways involved in lipid biosynthesis and storage, particularly sphingolipids, were consistently downregulated in CU x PR when compared to both of these cohorts as were those related to pathways related to tissue development and cell proliferation (“regulation of epithelial to mesenchymal transition”, “regulation of cell cycle process”).

**Figure 4.**
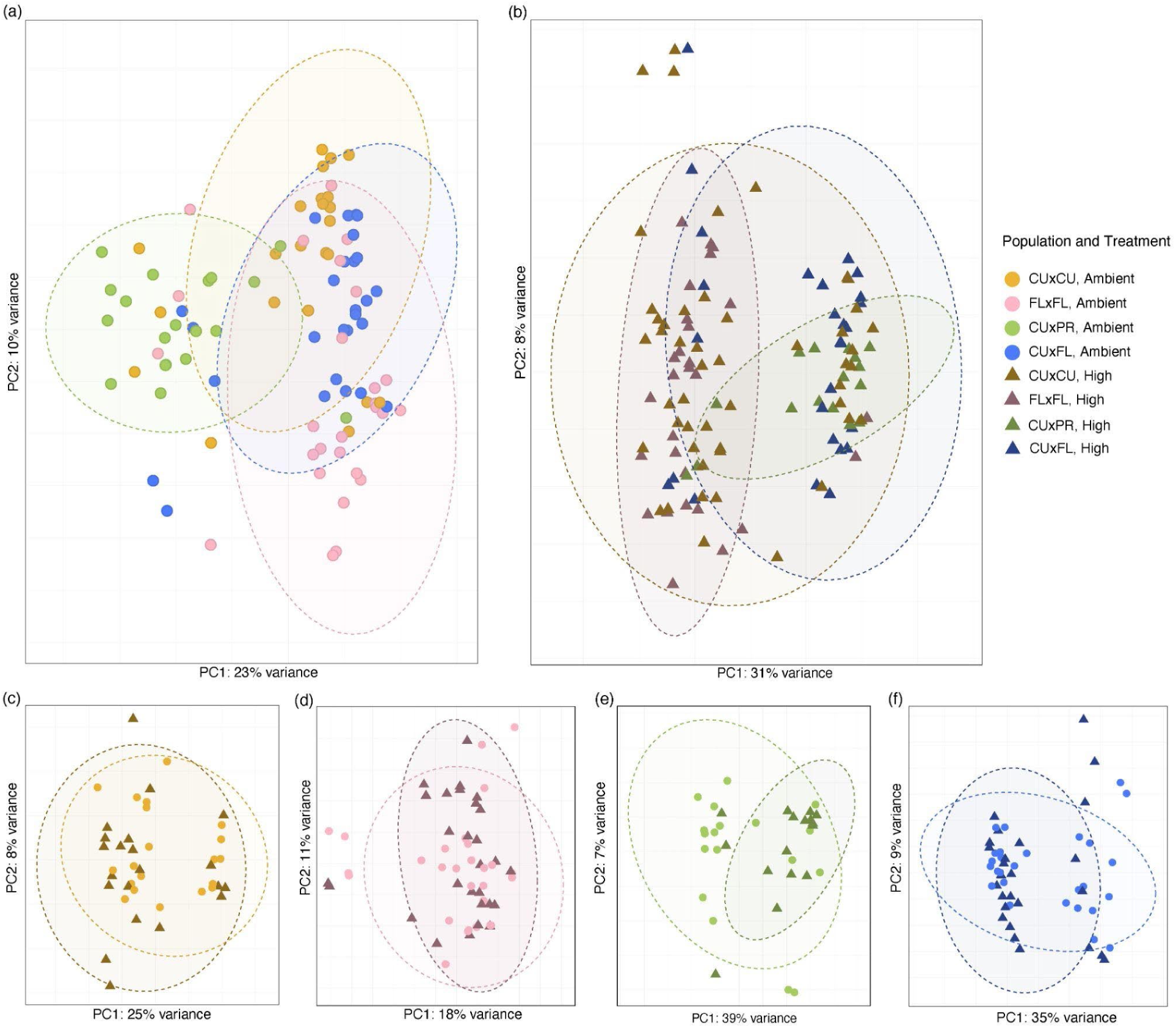
Principal Component Analysis (PCA) of gene count data across cohorts and temperature treatment (ambient temperature samples denoted by lighter colored circles and elevated temperature samples shown as dark-shade triangles). Baseline gene expression profiles of the four cohorts (*a*) primarily separates the Curaçao x Puerto Rico (CU x PR, green) AGF crosses from the remaining crosses, explaining 23% of the variance between samples along Principal Component 1 (PC1), while 10% of variance explained by PC2 appears to be driven by the transcriptomic divergence between the non-AGF Curaçao x Curaçao (CU x CU, yellow) and Florida x Florida (FL x FL, pink) genets. After elevated temperature exposure, temperature treatment did not drive population differentiation, with the four cross cohorts clustering indiscriminately together (*b*); none of the variables tested explained the 31% of variance between populations captured along PC1. Neither CU x CU (*c*) nor FL x FL (*d*) cohorts showed a significant thermal stress response, only differentially expressing 34 and 20 genes, respectively. AGF cohort CU x PR (*e*) was more affected by the temperature change, with 175 DEGs explaining 39% of variance on PC1. Curaçao x Florida (CU x FL, blue, *f*) had 81 DEGs in response to thermal stress, though the 35% of variation explained by PC1 is not driven by this. Clustering patterns seen in the CU x CU and CU x FL are instead likely driven by genet-specific variation rather than treatment response.

Curaçao (CU x CU) crosses also showed significant transcriptomic differences, with 2,841 DEGs when compared to CU x PR. While less extreme than the CU x PR comparisons, the CU x CU cohort similarly diverged from Florida-containing cohorts, with approximately 180 DEGs for both CU x FL and FL x FL comparisons, leading to a slight separation in the PCA clustering of the CU x CU and FL x FL cohorts that explained 10% of variation along the second principal component (PC2; Figure 4*a*). In contrast, CU x FL and FL x FL demonstrated the highest transcriptomic similarity, with only 8 DEGs between them, indicating a highly conserved expression profile within Florida-derived lineages. Of these 8 differentially expressed genes, higher expression was found in the CU x FL cohort for the *collagen triple helix repeat-containing protein 1* (*cthrc1)* gene, a secreted glycoprotein potentially involved in regulating tip growth in *Acropora* (Hemond et al., 2014), as well as enrichment pathways related to wnt protein binding. The innate immune response gene *C-type mannose receptor 2* was upregulated however in the FL x FL cohort.

### Transcriptomic Response to Elevated Temperature Conditions, Within Populations

Of the four cohorts, only the hybrids CU x PR and CU x FL showed a significant transcriptomic change at the end of the experiment after thermal stress exposure, with 175 and 81 differentially expressed genes found (respectively), compared to the 34 and 20 DEGs in the CU x CU and FL x FL cohorts. PCA analyses subsequently showed substantially overlapping distributions of treated and ambient samples for these latter two cohorts (Figure 4*c, d*), with the two somewhat separated clusters seen in the CUxCU cohort more likely explained by greater genotypic diversity between individuals. In contrast, samples separated based on treatment into two clusters for both the CU x PR cohort (Figure 4*e*) and CU x FL cohort (Figure 4*f*), albeit weakly.

The most studied candidate genes in coral related to stress response are those encoding heat shock proteins (HSPs), with *hsp70* often upregulated during high temperatures. Only found in the CU x PR cohort, *hsp70* was instead under-expressed (-0.58 log2-fold decrease) as were other stress response genes including *cytochrome P 450 3A18*, an oxidative stress related gene (-0.94 log2-fold change), an apoptosis regulator *TNF receptor-associated factor 6* (-0.89 log2-fold change), and *complement factor B,* a key player in innate immunity (-1.5 log2-fold change).

Despite 81 genes being differentially expressed, the CUxFL cohort’s transcriptional response at the conclusion of the experiment was mostly devoid of typical coral heat stress biomarkers except for the calcium-modulated protein, *Calmodulin*, and *Myosin-2 essential light chain*, both of which were slightly under-expressed. Notably absent was the upregulation of heat shock proteins and antioxidant genes commonly associated with thermal stress. Instead, the response was characterized by the upregulation of structural genes such as *collagen triple helix repeat-containing protein 1* and the downregulation of various metabolic and cellular process genes that are often over expressed and associated with oxidative stress responses (e.g., *universal stress protein*, *thioredoxin domain-containing protein*). Gene Ontology analysis revealed greater enrichment in terms related to tissue remodeling and morphogenesis, including “regulation of canonical Wnt signaling pathway” (GO:0060828) and “tissue remodeling” (GO:0048771).

Though showing minimal transcriptional response, the FL x FL cohort was the only other cohort that significantly upregulated any heat shock proteins, specifically *hsp16* (1.97 log2-fold increase), as well as enrichment pathways related to ‘response to heat’ (GO:0009408) and ‘response to temperature stimulus’ (GO:0009266). However, of the remaining 19 DEGs, none of the 8 annotated genes had any significant role in heat stress response.

Notably, the majority of DEGs across cohorts were downregulated by the conclusion of the experiment, except for CU x CU, which displayed a unique pattern of predominantly upregulated genes (33 vs 1 DEG). One such gene included *Sacsin* (2.3 log2-fold increase), a co-chaperone for the heat-shock protein Hsp70, which was significantly up-regulated when exposed to elevated temperatures in other *Acropora* and *Pocillipora* (Fuller et al., 2020).

### Transcriptomic Response to Elevated Temperature Conditions, Between Populations

The magnitude of pairwise transcriptomic differences between cohorts decreased after heat stress (Figure 4*b*), though generally followed the same trends as observed at baseline conditions; most differences were found when comparing CU x PR to the non-AGF CU x CU cross (162 DEGs), which included few genes known for regulating thermal stress, but did include “oxidative stress-responsive serine-rich protein 1” (log2FoldChange = -1.06; padj = 0.05), “Forkhead box protein L1” (log2FoldChange = -1.67; padj = 0.024), and “metalloproteinase inhibitor 3” (log2FoldChange = -2.09; padj = 0.003) all of which have tentative roles in oxidative as well as general stress response. 222 Differentially expressed genes were found between CU x PR and the FL x FL cohort with several transcripts showing notable upregulation. For example, ‘bacterial dynamin-like protein’ (log2FoldChange = 3.86; padj = 0.027) and ‘allene oxide synthase-lipoxygenase protein’ (log2FoldChange = 2.54; padj = 0.049) were the most over expressed genes in the CU x PR population corals compared to FL x FL, and transcripts commonly associated with stress responses were also differentially expressed. Of particular note, heat shock protein ‘Hsp-16.2’ (log2FoldChange = -1.75; padj = 0.041) was downregulated in CU×PR relative to FL×FL corals, while ‘peroxidasin’ was upregulated (log2FoldChange = 1.1, padj = 0.01).

The other hybrid population, CU x FL had a more similar response to the non-AGF populations, with 61 differentially expressed genes compared to the CU x CU population and only 8 differentially expressed genes when compared to FL x FL (with only 4 genes annotated, none of which have a known role in thermal stress response). When comparing CU x FL to CU x CU expression profiles, “TRAF6”, which mediates cellular stress signaling and immune responses, was significantly downregulated (log2FC = -1.17, padj = 0.041). “Cytochrome P450”, involved in oxidative stress responses, also showed decreased expression (log2FC = -1.79, padj = 0.013). Notably absent from the differentially expressed genes were heat shock proteins and other canonical thermal stress response genes.

Surprisingly, the hybrid cohorts CU x PR and CU x FL, which both had the greatest transcriptomic shifts (above) by the end of the experiment showed only 2 differentially expressed genes between them, neither of which are annotated, indicating a mostly conserved response to elevated temperatures.

CU x CU and FL x FL had similar transcriptomic profiles with only 15 differentially expressed genes between them after heat stress exposure.

To explore mid-parent deviation of gene expression in a hybrid population (Figure 5), normalized counts of the CU x FL population were compared to the mean of purebred FL x FL and CU x CU populations, yielding 26 upregulated and 48 downregulated genes in the hybrid cross (alpha=0.05, lfcThreshold=0.1). Pairwise comparisons show a greater similarity between the CU x FL and FL x FL cohorts (with 1 DEG, versus 20 DEG with cohort CU x CU).

**Figure 5.**
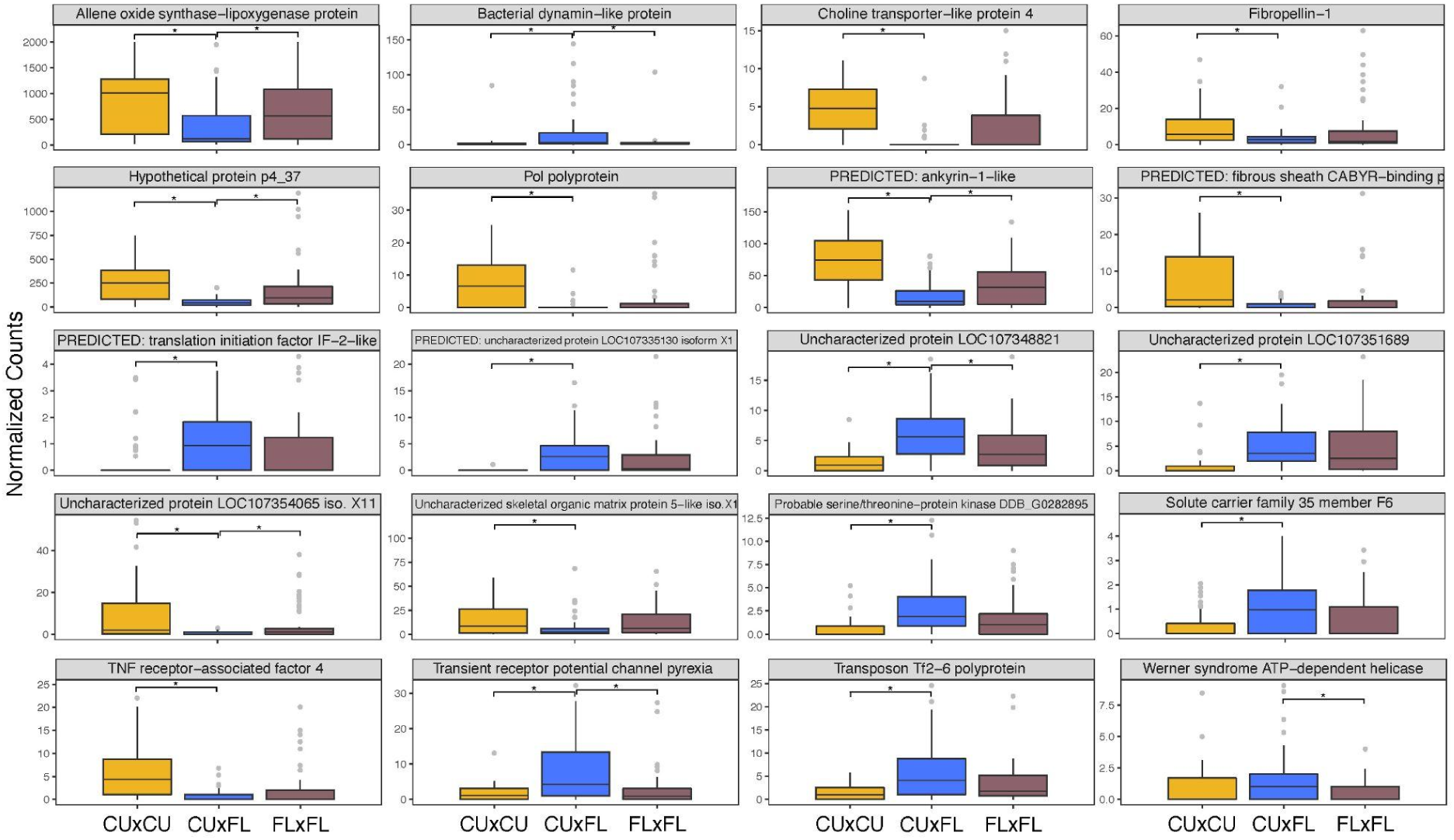
Expression differences in hybrid CU x FL cohort compared to pure parental cohorts. Boxplots show normalized gene counts for the top 20 most differentially expressed genes between the purebred parental cohorts FL x FL (brown), CU x CU (yellow), and hybrid cohort CU x FL (blue) under pre-exposure conditions

### Symbiont Typing

The results of both a linear discriminant analysis of the signal intensities and a second random forest analysis indicate that all corals used in the experiment were hosting predominantly algae in the genus *Durusdinium* (Supplementary Table 1).

### Association Analysis

Among all 19,695 SNPs, 6,700 polymorphic SNPs were retained for the association analysis. Across all 5 traits tested, no significant associations were detected. The trait that had the strongest overall contribution to the variation in the data (F_v_/F_m_) was selected to plot the distribution of p-values across chromosomes of the *Acropora palmata* genome (Figure 6).

**Figure 6:**
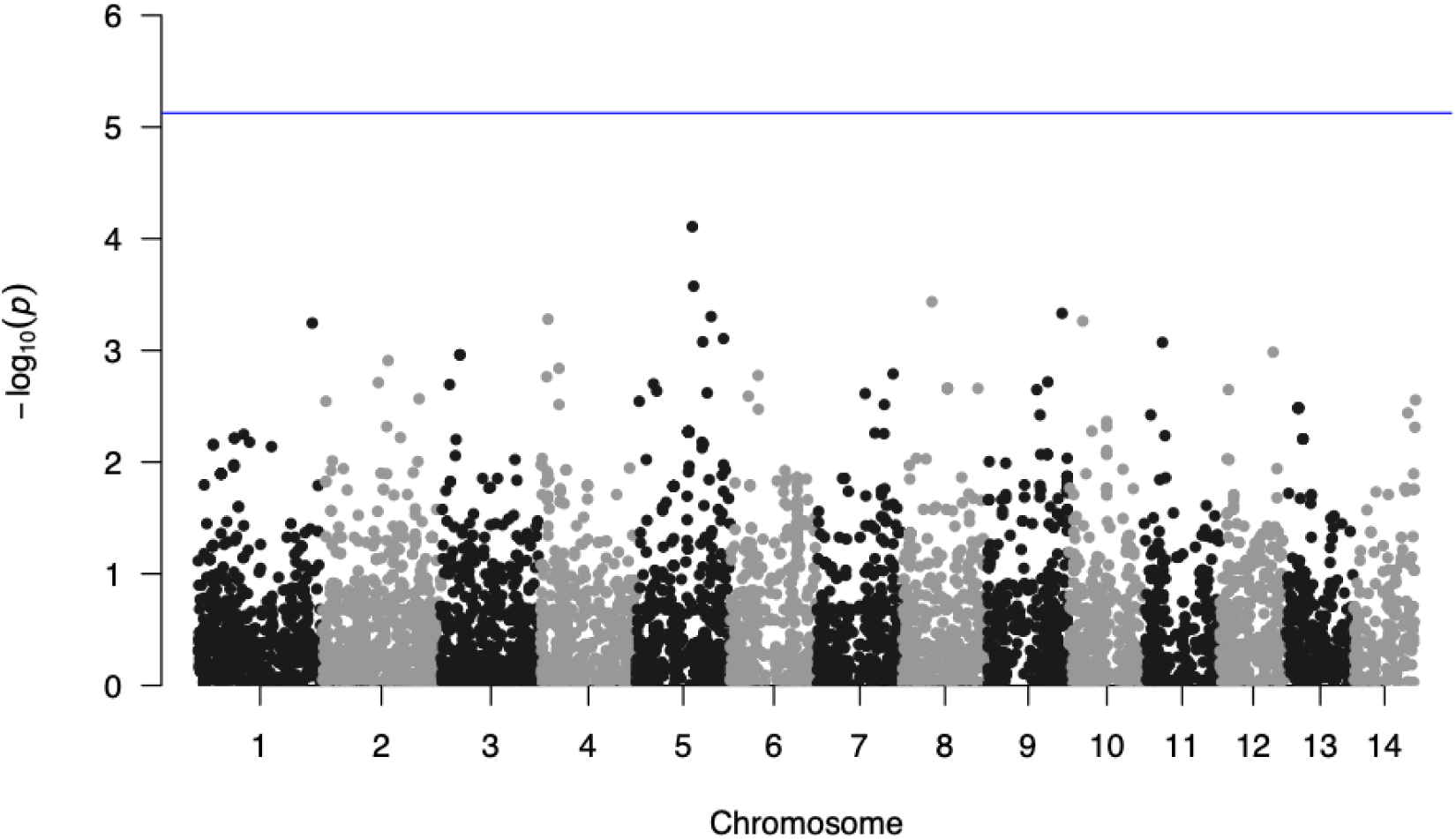
Manhattan Plot of Genome-wide p-values for SNPs associated with variation in Fv/Fm in cohorts tested in this study. Resolved chromosomes in the *A.palmata* genome along the x-axis (Locatelli et al., 2024). P-values of SNPs calculated using a linear model in plink2. Blue line represents the *p* at which a SNP would be considered significant after applying a Bonferroni correction. SNPs with a p-value of NA have been removed from analysis.

### Genetic Diversity

The principal component analysis (Figure 7a) shows that the Florida cohort is quite distinct from the other three cohorts. This was expected, as all three other cohorts contain some proportion of Curaçao ancestry. As well, the CU x FL cohort clustered more closely to the FL x FL cohort than any other hybrid cohort. The FL x FL cohort, while being genetically distinct from the three other cohorts, also has the largest genetic variation within itself, spreading across principal component 2. This is likely because corals in the FL x FL cohort were derived from a larger number of parents. In the CU x CU cohort, two subclusters are visible in the PCA (Figure 7a) but genetic variation with the CU x CU cohort is small due to the limited number of parents. When randomly selected wild samples are included in the principal component analysis (Figure 7b), the FL x FL cohort clusters well with the wild FL samples, while the AGF cohorts with Curaçao ancestry cluster much closer to each other rather than with the wild samples of their source populations, highlighting their unique genetic background.

**Figure 7:**
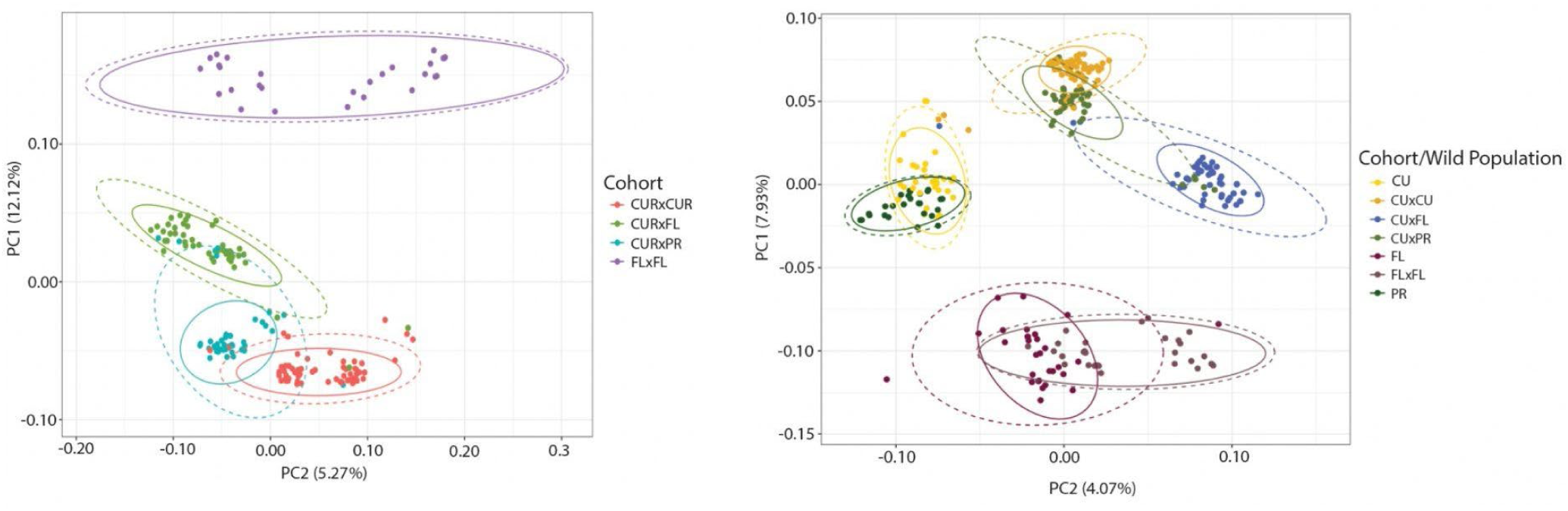
Genetic divergence of cohorts. a) Principal Component Analysis (PCA) of pairwise genetic differences between all samples calculated using 19,695 SNPs. 12.41% of variation is explained by Principal component (PC) 1 and 5.23% of variation is explained by Principal component (PC2) 2. As expected, the FLxFL corals cluster distinctly from the remainder of the cohorts, demonstrating their unique ancestry relative to the other samples. b) PCA of pairwise genetic differences of the four AGF cohorts and randomly selected wild genets from the three source populations (FL, CU, PR). FL wild and experimental cohorts are well differentiated from the other cohorts and populations while PR and CU wild populations are more genetically similar. CU x CU and the FL x FL crosses show shifts in allele frequencies relative to the wild population, likely due to stochasticity as well as selection imposed during larval rearing.

There were significant differences (p<0.05) in mean allelic diversity between all cohorts (Figure 8a). While the mean allelic richness was significantly different between the CU x CU and CU x PR cohorts, CU x FL and FL x FL cohorts (Figure 8a), it is important to note that the range of mean allelic richness of all cohorts was small [1.236 (CU x CU) to 1.299 (FL x FL)]. Between CU and FL wild populations, there were 2,963 private alleles, 2,026 of which were private to FL and 937 of which were private to CU. The CU x FL hybrid cohort contained 879 of private CU alleles and 1,984 of private FL alleles (Figure 7b). Between CU and PR wild populations, there were 2,166 private alleles, 922 of which were private to PR and 1244 of which were private to CU. The CU x PR cohort contained 1,153 private CU alleles and 877 private PR alleles (Figure 8c).

**Figure 8:**
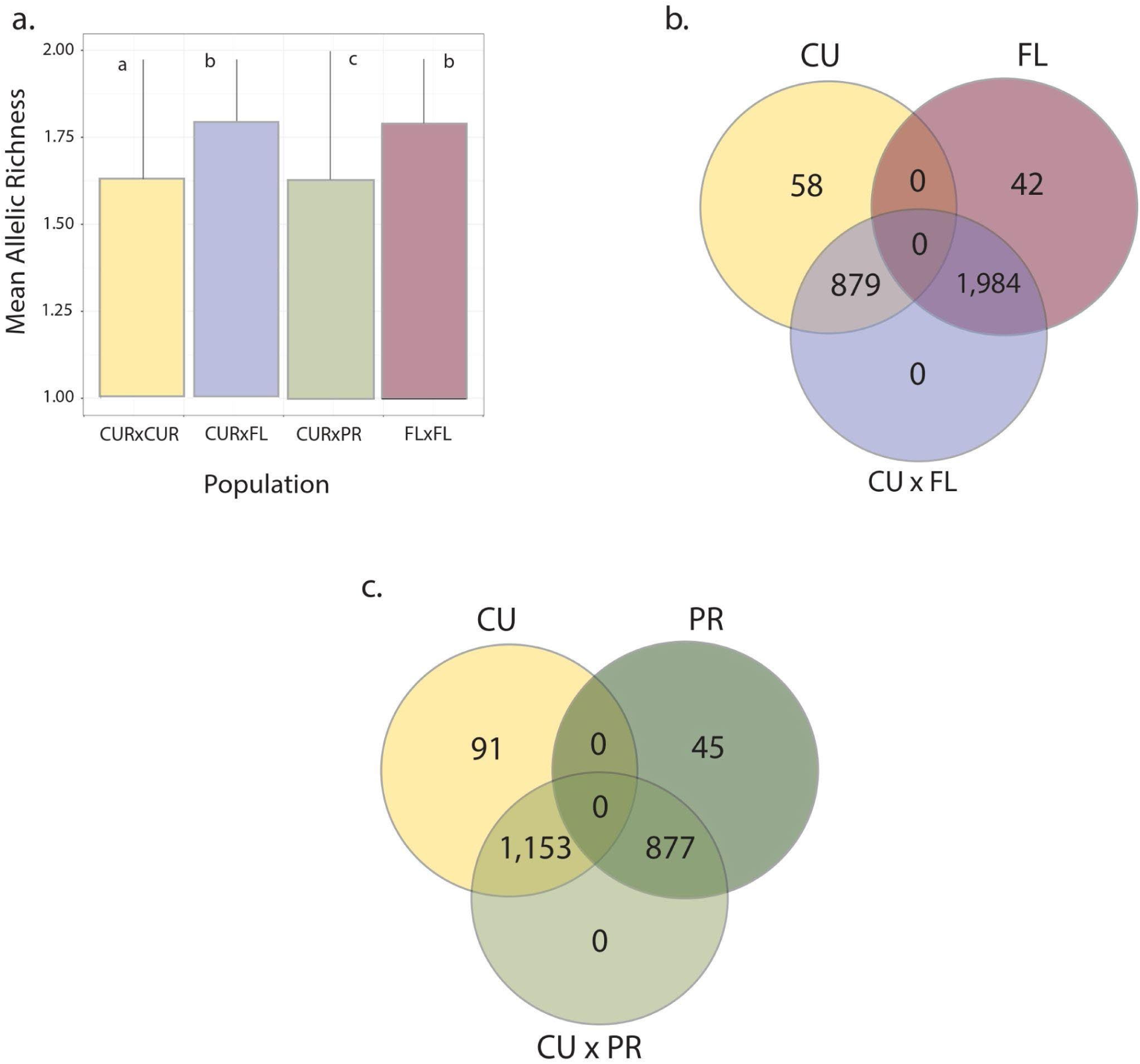
**Allelic diversity and private alleles in hybrid populations**. a) Mean allelic richness across all 19,695 loci. There were significant differences (indicated by lower letters) in allelic diversity across most cohorts. The FL x FL cohort shows the greatest mean allelic richness. However, there was little reduction in genetic diversity due to the smaller number of parents used to create the cohorts of Curaçao ancestry b) Venn Diagram of private alleles shared in hybrid CU x FL and c) CU x PR cohorts. The CU x FL cohort contained 879 alleles from Curaçao not previously recovered in the Florida populations, and the CU x PR cohort contained 1,153 alleles from Curaçao not previously recovered in the Western Puerto Rico populations.

Pairwise kinship coefficients were calculated between all corals in the experiment and analyzed within and between cohorts with an ANOVA. A mean kinship above 0 indicates that recruits are highly related. The CU x FL cohort had the highest mean kinship of 0.118 (second degree relationship) within its cohort, while the FL x FL population had the lowest mean kinship within its cohort of -0.0360 (unrelated). Overall, the FL x FL cohort contained more unrelated recruits. This reflects the results from the PCA, showing that the FL x FL cohort is not only genetically distinct, but also highly unrelated within itself due to the larger number of parents contributing.

Average observed heterozygosity of the CU x CU cohort was 0.136± 0.022. Average observed heterozygosity of the CU x FL cohort 0.135 ± 0.021. The average heterozygosity of the FL x FL cohort was 0.149. ± 0.024. The average heterozygosity of the CU x PR cohort was 0.138. ± 0.022. None of the cohorts significantly differed from expected heterozygosity. The CU x CU and CU x FL cohorts had slightly elevated FIS, while the CU x PR and FL x FL cohorts had slightly depressed FIS (Table 3).

**Table 3:** Heterozygosity of cohorts measured in the follow-up heat stress study. Observed and expected heterozygosity were calculated for each cohort examined in this study, along with FIS (inbreeding coefficient). No significant deviation from Hardy-Weinberg expectations were found.

| Cohort | N_Loci | Observed Heterozygosity | Expected Heterozygosity | FIS |
| --- | --- | --- | --- | --- |
| CU x CU | 19695 | 0.136 | 0.138 | 0.0098 |
| CU x FL | 19695 | 0.135 | 0.147 | 0.0797 |
| CU x PR | 19695 | 0.139 | 0.137 | 0.0126 |
| FL x FL | 19695 | 0.15 | 0.14 | 0.0726 |

## Discussion

In this study we present the first comparative dataset characterizing the genetic diversity and physiological responses to heat stress of F1 crosses among purebred and AGF cohorts of the endangered reef-building coral, *Acropora palmata*.

### High temperature exposure may not have reached the corals’ thermal threshold

Although the cohorts were exposed to water temperatures that caused extensive bleaching of *A. palmata* under natural reef conditions of the Florida Keys several times in the past decade (Manzello et al. 2007, Williams et al. 2017), the corals within the present experiment showed very few signs of physiological stress associated with exposure to 31 °C for two months. All cohorts even showed a trend of higher growth rates (2DSA, 3DSA, and BW) under the high temperature treatment, some of which were statistically significant (e.g., CU x PR 2DSA). Previous studies suggest a Gaussian distribution occurs between water temperatures and calcification rates where a positive correlation takes place until a thermal threshold is reached, the symbiosis between the coral host and the algal symbiont breaks down, bleaching occurs, and growth rates decline (Jokiel *et al*. 1978). Although growth rates were higher under high temperature conditions in the present study, these measurements represented an accumulation of the physiological response over the entire two-month time period, which may have largely captured conditions prior to the corals reaching their thermal exposure threshold. There was substantially less P_net_ occurring within the high temperature conditions of all cohorts measured at the end of the experimental treatments (after two months of exposure). End point measurements, such as P_net_, suggest physiological stress may have been occurring near the end of the experiment. Muller et al. 2021 suggests that end point measurements may be more sensitive phenotypic metrics compared with cumulative measurements, and González-Guerrero et al. 2021 show metrics such as gross photosynthesis may be more accurate than measurements acquired using PAM fluorometry. The results of the P_net_ and daily P:R ratios suggest that initial stages of a temperature threshold may have been reached after two months, but the experiment ended prior to the cohorts fully reaching the temperature stress threshold that would likely have resulted in greater physiological ramifications such as reduced growth.

### The FL x FL cohort was the most sensitive and the Curaçao cohorts were the most robust to elevated temperatures

Of all the cohorts tested, the FL x FL cross was the most sensitive to the high temperature exposure as it was the only one to show a significant reduction in P*_net_* and ETR*_max_* after two months of exposure to high temperatures. Interestingly, the CU x FL cohort showed a significant positive photochemical response associated with high temperature exposures with a greater F_v_/F_m_ value and a reduced RGB value (indicating higher algal symbiont densities) compared with the ambient conditions. Higher F_v_/F_m_ values were observed within the CU x CU cross under high temperature conditions as well. These initial results suggest that the hybrid cohort, which included both the Florida and Curaçao alleles, and the purebred CU x CU cohort may be more resistant than others, particularly compared with the purebred FL x FL cohort. Although preliminary, these are encouraging results that may be worth further exploring to assess whether AGF may be useful for climate-priming cohorts at risk to global warming.

### The dominant algal symbiont, Durusdinium trenchii, likely influenced phenotypic response

The lack of a strong negative stress response associated with the high temperature exposures from the cohorts may also be from the unique association between the *A. palmata* reared within the study and the dominant algal symbiont identified within each of the cohorts, *Durusdinium trenchii*. This particular algal symbiont is significantly more heat tolerant than other species of Symbiodiniacaea (LaJeunesse et al. 2014, Manzello et al. 2019), which may have provided thermal protection to the extended high temperature conditions within the study. Indeed, most *A. palmata in situ* are associated with a more temperature sensitive algal symbiont, *Symbiodinium* ‘*fitti’* (Baums et al. 2014, Durante et al. 2019). These results indicate that the experimental corals used within the present study may be more thermally tolerant than the naturally occurring populations because of the unique host/algal symbiont relationship, which developed as an artifact of rearing location, Mote’s land based coral nursery. Whether these more heat tolerant symbionts are retained after outplanting is not well known. Elder et al. (2023) showed that Mote-sourced corals outplanted with *D. trenchii* retained that algal symbiont for up to two years. These results suggested that the juvenile hosts and the novel symbiont may maintain their association for several years even after introduction to the natural reef environment even if the host eventually reverts to the analogous symbiont, *S. ‘fitti’*. On the contrary, Gantt et al. (2025) showed an almost immediate shuffling from *D. trenchii* to *S. fitti* of Mote-sourced corals, evident within just one year after outplanting. Indeed, further study is needed to fully understand the influence of this novel host-symbiont association and the persistence of the association, or not, through time.

### CU x FL hybrids contain novel allelic diversity not known to be present in the FL population

One of the benefits of AGF is the introduction of novel alleles to a declining population (Frankham et al., 2017). Standing genetic variation is important to provide the material for future adaptation (Barrett & Schluter, 2008). Florida acroporids have suffered catastrophic declines and only 158 genets are currently surviving in the wild (Williams et al. 2024). The AGF hybrids carry >1000 alleles that are not currently known to be present in FL population according to assay of 19,696 loci, and as such present a valuable source of novel alleles and means to increase standing genetic variation within Florida which is not achievable by pure-bred crosses. The genotyping array indicated that 879 novel alleles could be introduced from Curaçao into the Florida population based on the 19,696 SNP loci assayed. The CU x FL AGF cohort thus may provide substantial additional genetic diversity to avoid inbreeding and allow a declining FL population to adapt to changing environmental conditions. While the Curaçao wild population is much more genetically similar to Western Puerto Rico (Figure 7a.) compared to the other populations, there were still 1,153 private Curaçao loci that could be potentially introduced as adaptive standing genetic variation into Puerto Rico populations.

The genetic data also highlighted that coral cohorts created from a limited number of parents are necessarily closely related. It is therefore advisable that outplanting of offspring from the same cohort is planned so that the risk of future inbreeding is considered. This is best achieved by limiting the number of related recruits that are placed in close proximity to each other on a reef. Recent data from the Pacific (Mumby et al. 2024) indicated that in that system, a distance of over 10 m should be effective in severely reducing fertilization opportunities.

### AGF cohorts provide novel differential gene expressions and parallel phenotype

Within-species, between-population hybrid gene expression analyses on corals have rarely been done before on corals. Crosses within and between *Acropora millepora* parents from a high and a low latitude population revealed that molecular and phenotypic thermo-tolerance has a heritable component (Dixon et al. 2015). Similar results were obtained when crossing *Platygyra daedalea* mothers from the relatively cool regions in the Indian Ocean with fathers from hot reefs in the Persian Gulf (Howells et a. 2021). Hybrid cohorts had a 37% increase in heat survival over purebreds from the cold Indian Ocean location. Both studies report sometimes large differences in stress tolerance by family, regardless of origin. Stress tolerance in *P. daedalea* could be associated with fathers carrying certain beneficial alleles, regardless of their population origin. In line with these results, we report here that the hybrid crosses deviated from the mid-parent value in tens of genes. Of the annotated genes, some were related to metabolic processes, growth, and innate immune regulation, as well as regulation of cell apoptosis. The hybrid CU x PR cohort had the greatest number of DEGs under baseline conditions and after stress treatment. The CU x FL hybrid cohort more closely resembled the FL x FL purebred cohort and was more dissimilar to the CU x CU purebred cohort. The gene expression patterns thus reflect the phenotypic results, indicating that the CU x CU cohorts were less impacted by the stress. Novel gene expression patterns by the hybrid AGF cohort CU x FL represent the conservation value of the AGF corals in introducing novel molecular phenotypes to Florida. AGF corals have not been outplanted into the wild. Should the novel alleles carried by AGF corals prove maladaptive to Florida Reefs, the likely outcome is failure of sexual reproduction and/or death of the colonies. Novel alleles may cause outbreeding depression in later generations, but this cannot be known until F2 or F3 corals exist. Given the lack of successful natural sexual reproduction in Florida (Williams et al. 2008) that risk seems quite low.

### Conclusion

This work demonstrates the potential power of a mixed provenance strategy when conducting active restoration for the purpose of species conservation. The perpetual poor performance of local populations in recent years, e.g., *A. palmata* in Florida’s Coral Reef, may suggest that whatever local adaptation is present in these populations is insufficient to ensure persistence as environmental conditions continue to change and decline (Williams *et al*. 2008; Williams *et al*. 2014; Baums *et al*. 2019; Chan *et al*. 2019). Thus, populations may be so degraded that the genetic risks associated with assisted gene flow (e.g., outbreeding depression) are nominal. The present study showed high levels of phenotypic similarity among the four cohorts tested even after exposure to elevated temperatures for two months and despite the varied number of parents contributing to each. The novel relationship between the coral host and *Durusdinium trenchii*, a resulting phenomenon associated with reared *Acropora* spp. within Mote’s land-based coral nursery, suggests all of these cohorts may be more resistant to thermal stress compared with those reared within the wild. Regardless, the native Florida cohort appeared slightly more sensitive to exposure to high temperatures whereas the AGFAGF CU x FL and the purebred purebred CU x CU cohorts were the most robust. The AGF cohorts were also more similar to the FL x FL purebred cohort in both phenotype characteristics and gene expression. Additionally, CU x FL individuals harbor genetic diversity not previously found in Florida that could be potentially adaptive. An assisted gene flow strategy utilizing distantly located populations along an environmental gradient can therefore promote the transfer of adaptive variation into genetically-depauperate populations as a means of genetic rescue. However, depending on the mechanism(s) underlying outbreeding depression, fitness reductions may be observed within F1 generations but perhaps not until after the F2 generation (Baums *et al*. 2008; Willis *et al*. 2006). Therefore, to understand the long-term impacts of AGF on coral fitness, further research is necessary, which could include controlled breeding studies and common garden experiments.

## Acknowledgements

This work was supported by the Paul G. Allen Family Foundation (Request #12894). We would like to thank Mary Hagedorn, Christopher A. Page, Daisy M. Flores, Lucas Tichy, Valerie F. Chamberland, Claire Lager, Nikolas Zuchowicz, Kathryn Lohr, Harvey Blackburn, Tali Vardi, Jennifer Moore, Mark J. A. Vermeij, and Kristen L. Marhaver for creating these AGF cohorts (See Hagedorn et al 2021 for permits). We would like to thank Austyn Bushman, Jacob Harpring, Emily Williams, Rachel Serafin and Jenny Lee for assisting with coral husbandry and data collection at The Florida Aquarium. We also thank Sophie Wong, Ciara Smith, Suni Chutkan, Chloe Spring, Omar E. Elliott, and Danielle Atkins for assisting with coral husbandry, coral health assessments, and data collection as part of the physiological assays. These individuals, in addition to the following, also assisted with sample collection: Allison Delashmit, Grace Klinges, Christina Giralt, Kyle Knoblock, Curtie Leopold, Yuen Azu, Elyse Anderson, Chloe Spring, and Ian Combs. We thank Dana Williams and Katie Flynn for making SNPchip data available via the STAGdb database. Sheila Kitchen made available the *A. palmata* genome. While working on this project, EMM was supported by a Mote Eminent Scholarship, CCO was supported by an NSF-GRFP (#2020289929), and TC was supported by a PSU Eberly College SAGF fellowship.

## Supplemental Documents

**Supplementary Table 1**: Detected Symbiodiniaceae in *A. palmata* tissue samples. The raw signal intensities for each sample (in rows) and probe (columns D - AM) are given. Each sample is identified by the automatically generated sample id (Column A), the user specimen ID (Column B), and the cohort (Column C). The first five linear discriminants of the linear discriminant analysis are listed in Columns AN - AR. Their combination yields the assignment of Symbiodiniacea to clades (Columns AS - AX) following Kitchen et al., 2020.

